# Optimizing drug infusion schedules towards personalized cancer chronotherapy

**DOI:** 10.1101/688606

**Authors:** Roger J. W. Hill, Pasquale F. Innominato, Francis Lévi, Annabelle Ballesta

## Abstract

**Aims:** Precision medicine requires accurate technologies for drug administration and proper systems pharmacology approaches for patient data analysis. Here, plasma pharmacokinetics (PK) data of the OPTILIV trial in which cancer patients received oxaliplatin, 5-fluorouracil and irinotecan via chronomodulated schedules delivered by an infusion pump into the hepatic artery were mathematically investigated.

**Methods:** A pump-to-patient model was designed in order to accurately represent the drug solution dynamics from the pump to the patient blood. It was connected to semi-mechanistic PK models to analyse inter-patient variability in PK parameters.

**Results:** Large time delays of up to 1h41 between the actual pump start and the time of drug detection in patient blood was predicted by the model and confirmed by PK data. Sudden delivery spike in the patient artery due to glucose rinse after drug administration accounted for up to 10.7% of the total drug dose. New model-guided delivery profiles were designed to precisely lead to the drug exposure intended by clinicians. Next, the complete mathematical framework achieved a very good fit to individual time-concentration PK profiles and concluded that inter-subject differences in PK parameters was the lowest for irinotecan, intermediate for oxaliplatin and the largest for 5-fluorouracil. Clustering patients according to their PK parameter values revealed two patient subgroups for each drug in which inter-patient variability was largely decreased compared to that in the total population.

**Conclusions:** This study provides a complete mathematical framework to optimize drug infusion pumps and inform on inter-patient PK variability, a step towards precise and personalized cancer chronotherapy.

**Author summary:** Accuracy and safety of infusion pumps remain a critical issue in the clinics and the development of accurate mathematical models to optimize drug administration though such devices has a key part to play in the advancement of precision medicine. Here, PK data from cancer patient receiving irinotecan, oxaliplatin and 5-fluorouracil into the hepatic artery via an infusion pump was mathematically investigated. A pump-to-patient model was designed and revealed significant inconsistencies between intended drug profiles and actual plasma concentrations. This mathematical model was then used to suggest improved profiles in order to minimise error and optimise delivery. Physiologically-based PK models of the three drugs were then linked to the pump-to-patient model. The whole framework achieved a very good fit to data and allowed quantifying inter-patient variability in PK parameters and linking them to potential clinical biomarkers via patient clustering. The developed methodology improves our understanding of patient-specific drug pharmacokinetics towards personalized drug administration.

## Introduction

Cancer management is challenged by large inter- and intra-patient variabilities in both disease progression and response to treatments. Thus, the quest for accurate and personalized cancer therapies has fostered the development of new technologies enabling multi-type measurements in individual patients and complex drug scheduling. To translate datasets available for an individual patient into personalized therapies and further ensure their precise administration, new mathematical approaches are required. Indeed, systems medicine, that involves the implementation of theoretical approaches in medical research and practice, is critically needed as emphasized in the roadmaps of the Coordinated Action for Systems Medicine (CaSyM) from the European Union (https://www.casym.eu, [1]) and of the Avicenna action (http://avicenna-isct.org/), and in other international consortia [2–5]. The final aim is a measurable improvement of patient health through systems-based practice which will enable predictive, personalised, participatory and preventive (P4) medicine [6].

Accuracy and safety of infusion pumps are mandatory to ensure that the correct drug dose is delivered to the patient over the intended period. Recurrent incidents related to devices delivering fluids such as nutrients or medications into the body have led the U.S Food and Drug Administration (FDA) to launch in 2010 an initiative to reduce infusion pump risks (https://www.fda.gov/medicaldevices/productsandmedicalprocedures/generalhospitaldevicesandsupplies/infusionpumps/ucm202501.htm). Many of the reported events are related to deficiencies in the initial design of the device and of the embedded software. Adverse events may also arise from a defect appearing over the device’s life cycle due to technical failure or lack of proper maintenance. However, due to the complexity of the equipment, user errors are also common [7].

Optimizing chemotherapeutics index, defined as the ratio between treatment antitumor efficacy and induced toxicities, is complex at multiple levels. First, large inter-patient variabilities are demonstrated in drug pharmacokinetics, tolerability and anti-tumour efficacy [2, 8–10]. Next, important intra-patient variabilities arise from the fact that tumour and healthy tissues, rather than being static over time, display time-dependent variations, in particular over the 24h span, which are called circadian rhythms [11]. The circadian timing system controls most physiological functions of the organism resulting in drug Absorption, Distribution, Metabolism and Elimination (ADME) displaying 24h-rhythms with differences of up to several folds between minimum and maximum activities [12, 13].

Chronotherapy -that is administering drugs according to the patient’s biological rhythms over 24 h- is a growing field in medicine and especially in oncology. Indeed, at least 22 clinical trials involving a total of 1773 patients with different types of metastatic cancers have demonstrated a significant influence of administration timing on the tolerability of 11 commonly-used antitumor drugs [14]. Two randomized phase III clinical trials in 278 metastatic colorectal cancer (mCRC) patients receiving irinotecan, oxaliplatin and 5-fluorouracil showed that cancer chronotherapy achieved an up-to-5-fold decrease in treatment side effects and nearly doubled anti-tumour efficacy compared to conventional administration of the same drug doses [15]. However, a meta-analysis of these two studies combined to another clinical trial involving 564 mCRC patients receiving the same drugs (497 men and 345 women in total) concluded that the chronomodulated drug modality significantly increased the efficacy and survival in men while reducing that in women as compared to conventional administration [16]. Such sex-specificity was further validated for irinotecan chronotoxicity in mouse experiments [17] and in a clinical trial involving 199 mCRC patients treated with oxaliplatin (infusion peak 4pm), 5-fluorouracil (infusion peak 4am) and irinotecan given at 6 different circadian times (https://academic.oup.com/annonc/article/28/suppl_5/mdx393.048/4109820). Both studies showed a higher circadian amplitude in females as compared to males and a difference of several hours between the optimal timing of each gender. Furthermore, circadian biomarker monitoring in individual patients recently revealed up to 12 h inter-patient differences regarding the timing of midsleep, the circadian maximum in skin surface temperature or that in physical activity [18]. These investigations have highlighted the need for the individualization of drug combinations and chronoinfusion schemes to further improve treatment outcome, taking into account the patients’ sex, chronotype and genetic background. The accurate delivery of intended administration profiles is obviously critical in this context. Chronotherapy requires the error in drug infusion timing not to be greater than few minutes.

Clinical findings about cancer chronotherapy have motivated the development of innovative technologies for chronomodulated drug delivery including the Mélodie infusion pump (Axoncable, Montmirail, France, [19]). This portable electronic pump allows for the administration of up to 4 compounds according to pre-programmed schedules over the 24 h span. It was used in several clinical trials for the chronomodulated delivery of irinotecan (CPT11), oxaliplatin (L-OHP) and 5-fluorouracil (5-FU) into the central vein of metastatic colorectal cancer patients [13]. The Mélodie pump was recently used to infuse those three anticancer drugs directly into the hepatic artery of metastatic cancer patients in the translational European OPTILIV Study [19]. In this study, the plasma pharmacokinetics of oxaliplatin revealed inconsistencies between programmed delivery schedules and observed drug concentration within the patient blood including a delay in the time taken for the drug to be detectable in the blood and unexpected peaks in plasma concentrations during drug infusion. Such inconsistencies between targeted drug exposure patterns and plasma drug levels motivated the design of a mathematical model of fluid dynamics within the pump system presented hereafter. This pump-to-patient model was then connected to semi-physiological PK models to investigate the inter-patient variability in drug PK after hepatic artery administration. Thus, this systems pharmacology study aimed to develop predictive mathematical models allowing for the quantitative and general understanding of i) the pump dynamics, irrespective of the drug delivery device, and ii) patient-specific whole-body PK of irinotecan, oxaliplatin and 5-fluorouracil after drug administration using an infusion pump. Such mathematical techniques would then allow for precise and personalized drug timing.

## Results

The overall objective of this study was to accurately investigate the inter-patient variability in the plasma PK of the three anticancer drugs administered during the OPTILIV trial. A first strategy consisted in using compartmental PK modelling taking the delivery profiles programmed into the infusion pump as inputs for the plasma compartments. However, such methodology revealed inconsistencies between the best-fit models and the data, including delays of several hours. We then concluded that the fluid dynamics from the pump to the patient had to be quantitatively modelled. Hence, we designed the complete model in two sequential mathematical studies. First, we studied the drug solution dynamics from the pump to the patient blood for which the model was based on partial differential equations. This novel model of the pump delivery system took into account the specificity of the equipment used in order to accurately predict drug delivery in the patients’ blood, although it could be easily adapted to any drug delivery devices. Second, we connected this model to compartmental PK models based on ordinary differential equations. This complete framework allowed the investigation of inter-patient variability in drug PK after hepatic artery administration.

### Pump-to-patient drug solution dynamics

#### Model design

The pump-to-patient model is a transport equation representing the dynamics of the drug solution along the administration tube, with respect to time (*t*) and one-dimensional space (*x*)(equation 1). *x* is the distance along the tube from the pump (*x* = 0) to the patient (*x* = *L*). The drug solution was assumed to be incompressible so that the fluid velocity was considered as constant along the whole tube. Thus, the drug concentration in the tube *u*(*x, t*) changes with respect to the following equation:

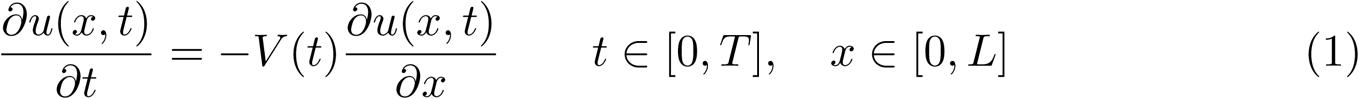

with a Dirichlet boundary condition of,

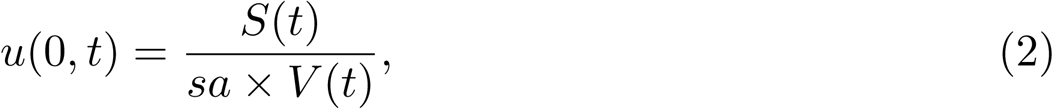

where *V* (*t*) is the fluid velocity inside the tube, expressed in mm/h. The constant *sa* = *πr*^2^ is the cross sectional surface area of the tube (in m^2^), with *r* being the radius of the tube. The source term *S*(*t*) represents the amount of drug delivered according to the infusion profile programmed into the pump and is expressed in mol/h. Initial conditions along the tube are *u*(*x*, 0) = 0. The fluid velocity and source terms are controlled by the pump which imposes a fluid delivery rate expressed in ml/h. They are computed by converting the fluid delivery rate into mm/h and mol/h respectively using the tube geometry and the concentration of each drug solution. Hence, model simulations at the end of the tube (*x* = *L*) do not depend on the exact geometry of the tube but rather on its total volume. The input function for PK models depending only on quantities at the end of the tube, the original infusion tube which was constituted of two sections of different diameters was simplified in numerical simulations to a tube of radius 1mm and total length 2340mm that had the same total volume as the original set-up. The total tube volume was set to 1.84 mL as in the equipment used in the OPTILIV study. The transport equation with associated initial and boundary conditions can be solved using the classical method of characteristics which gives [20]:

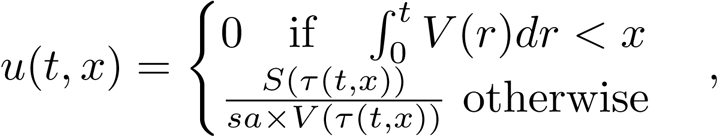

where *τ* (*x, t*) is the time at which the drug reaching point x at time *t* initially entered the system i.e.

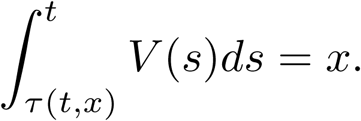

The input function for the PK models corresponds to the rate of drug infusion into the patient (i.e. at *x* = *L*) and can be obtained by:

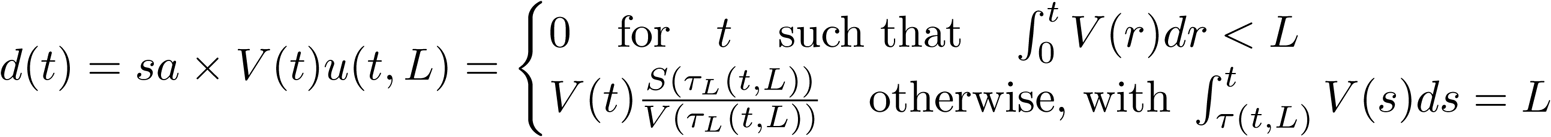

Note that, for all drug infusion apart from the glucose flushes, the source term S(t) is proportional to the fluid velocity *V* (*t*) as the drug is infused within the tube in the same time as the fluid, so that *d*(*t*) is proportional to *V* (*t*) once the tube is filled i.e. for times t such that 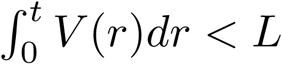. An example of the PDE model simulations in time and space for oxaliplatin delivery is shown in Fig 1a.

**Fig 1.**
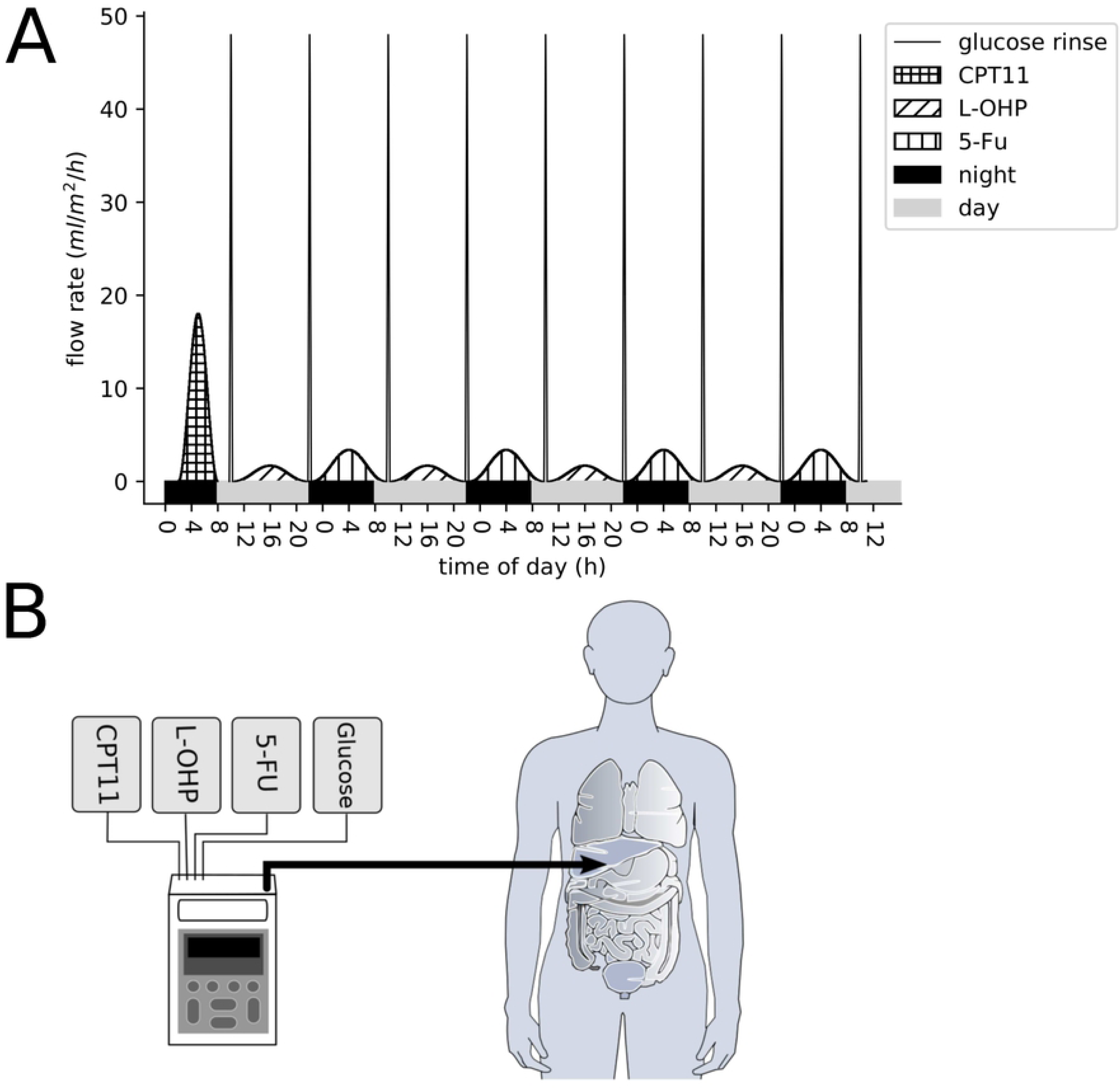
(a) shows oxaliplatin concentration profile in the infusion tube. The x-axis represents the distance along the tube, the y-axis represents the time from the start of the pump delivery. For figures (b-g), the x-axis represents Clock time and starts at the beginning of the considered drug administration. The left column shows the difference between the intended delivery profiles and the simulated delivery profiles evaluated at the end of the tube (x=L), for irinotecan (b), oxaliplatin (d) and 5-fluorouracil (f). The right-hand column shows the cumulative percentage of drug delivered to the patient for the intended and actual profiles over time for irinotecan (c), oxaliplatin (e) and 5-fluorouracil (g).

#### Differences between programmed infusion profiles and actual drug delivery in the patient’s blood

The pump infusion schemes used in the OPTILIV trial were simulated for the three drugs: irinotecan, oxaliplatin and 5-fluorouracil. Whereas the drug profiles programmed into the pump followed a smooth sinusoidal function, the actual drug delivery in the patient artery differed from the programmed profiles by two main features. First, the model predicted a significant time delay between the actual start of the drug delivery by the pump and the time the drug first reached the patient blood (Fig 1b,d,f). This delay was evaluated by the model to 3 h 5 min for oxaliplatin, 2 h 202 min for 5-fluorouracil and 51 min for irinotecan. It corresponded to the time taken to fill the infusion tube with the solution containing the drug at the beginning of the infusion. The delay was drug-specific as it depended on the drug solution concentration and the velocity of the solution in the tube driven by the programmed input profiles. Next, at the end of the infusion profiles, the pump stopped and did not administer the amount of drug left inside the tube. This remaining drug was flushed out by the glucose rinse subsequent to drug administration which induced a sudden delivery spike in the patient artery (Fig 1c,e,g). The amount of drug in this spike was expressed in percentage of total drug delivered and was estimated to 10.7% for oxaliplatin, 5.36% for 5-fluorouracil and 1.85% for irinotecan.

Our systems approach revealed important differences between the intended drug infusion profile and the actual administration into the patient artery. Hence, we developed optimized infusion profiles that strictly achieved the drug administration intended by clinicians. The same equipment was considered to avoid cost of changing. Drug concentrations of the infusion solutions were kept unchanged in order to avoid possible problems of drug stability. In order to administer the drug in the patient’s blood following a smooth sinusoidal function, a profile in three parts is required as follows (Fig 2). The first part of the profile is an initial bolus to fill the tube between the pump and the patient with the drug solution. Once the tube is filled, the original sinusoidal profile starts. Then, to solve the problem of the amount of drug left in the tube when the pump stops, the original sinusoidal profile needs to be interrupted when the total drug amount has left the drug bag. Then, a subsequent glucose rinse needs to be infused according to the final segment of the sinusoidal curve in order to deliver the drug remaining in the tube at the correct rate.

**Fig 2.**
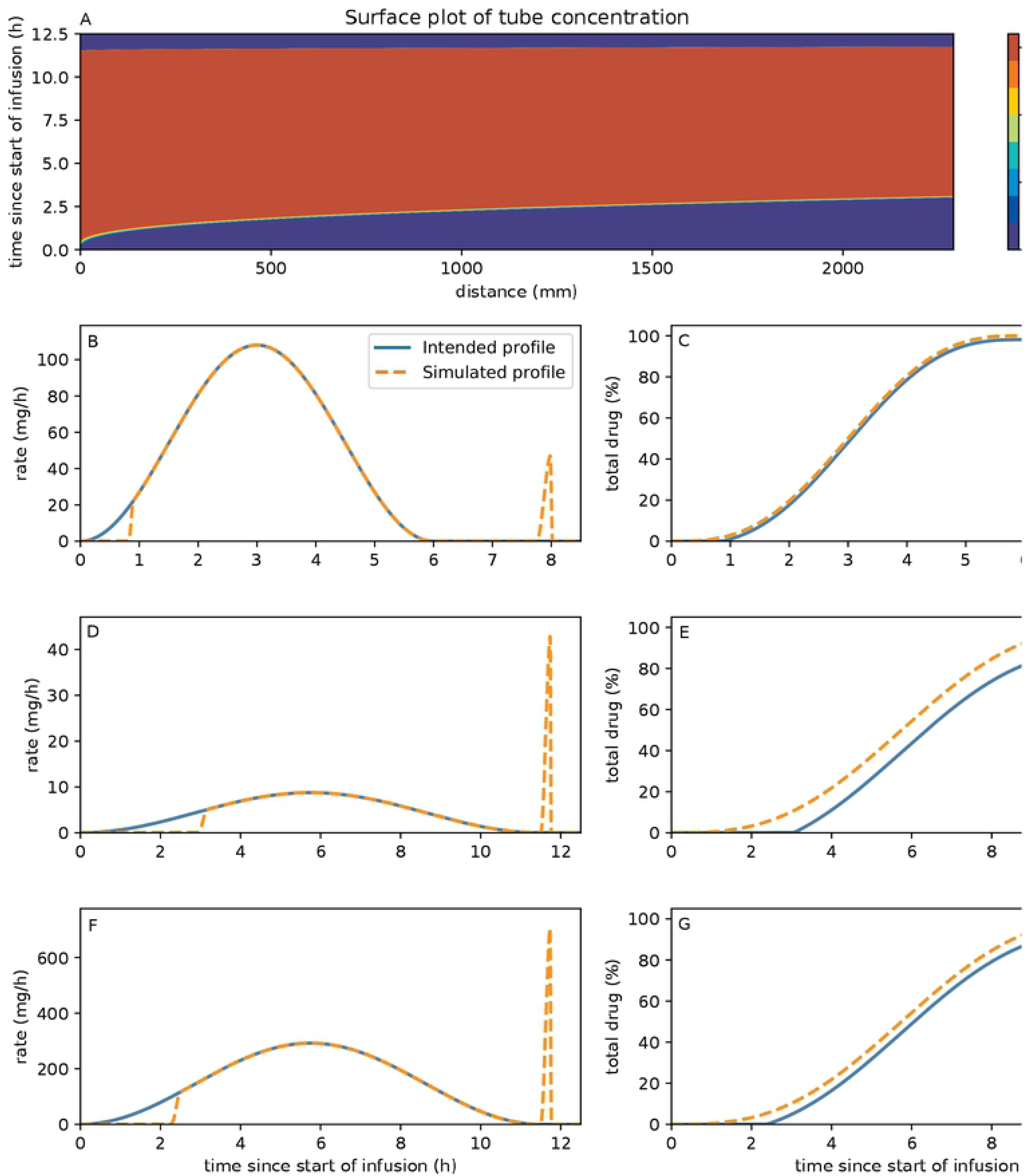
Improved administration profiles. (a) shows the drug solution delivery profile which consists of an initial bolus to fill the tube entirely, followed by the original profile. (b) shows the rinse solution delivery rate which continues drug delivery at correct rate while clearing the tube from any active substance, (c) shows how the flow rate along the tube is smoothly switched between the drug and the rinse and (d) shows the new drug delivery profile that will enter the patient compared to the original profile used in the OPTILIV study.

### Inter-patient variabilities in irinotecan, 5-fluorouracil and oxaliplatin PK after chronomodulated administration

The pump-to-patient model provided educated predictions of the drug infusion into the patients’ blood, which was a prerequisite to study the inter-patient variability in the PK of irinotecan, oxaliplatin and 5-fluorouracil. A compartmental physiological model was designed for each drug and all parameters were fitted for each patient independently.

#### Compartmental models of irinotecan, oxaliplatin and 5-fluorouracil pharmacokinetics

PK models represented the drug fate in: the Liver, to accurately represent hepatic delivery, the Blood, the measurement site, and the rest of the body known throughout this paper as Organs. The volume of each compartment was individualised for each patient using Vauthey method for Liver [21], Nadler’s formula for Blood [22], and Sendroy method for Organs [23]. Each model assumed that the drug was delivered directly into the liver compartment to represent the Hepatic Artery Infusion (HAI, Fig 3, 4 & 5). All transports in between compartments were considered as passive and represented by linear kinetics. Drug clearance included renal elimination for the Blood compartment, intestinal elimination for the Organs compartment and biliary excretion for the Liver compartment. In the absence of quantitative data and to avoid model over-parametrization, circadian rhythms were neglected in the PK models and all parameters were assumed to be constant over the 8-hour time window of PK measurements. Any chemical species bound either to plasma proteins or to DNA was assumed to be unable to move between compartments or to be cleared from the system. For the sake of simplicity, uptake and efflux rate constants were assumed to be equal for Blood-Liver and Blood-Organs transport respectively, for each of the three drugs.

**Fig 3.**
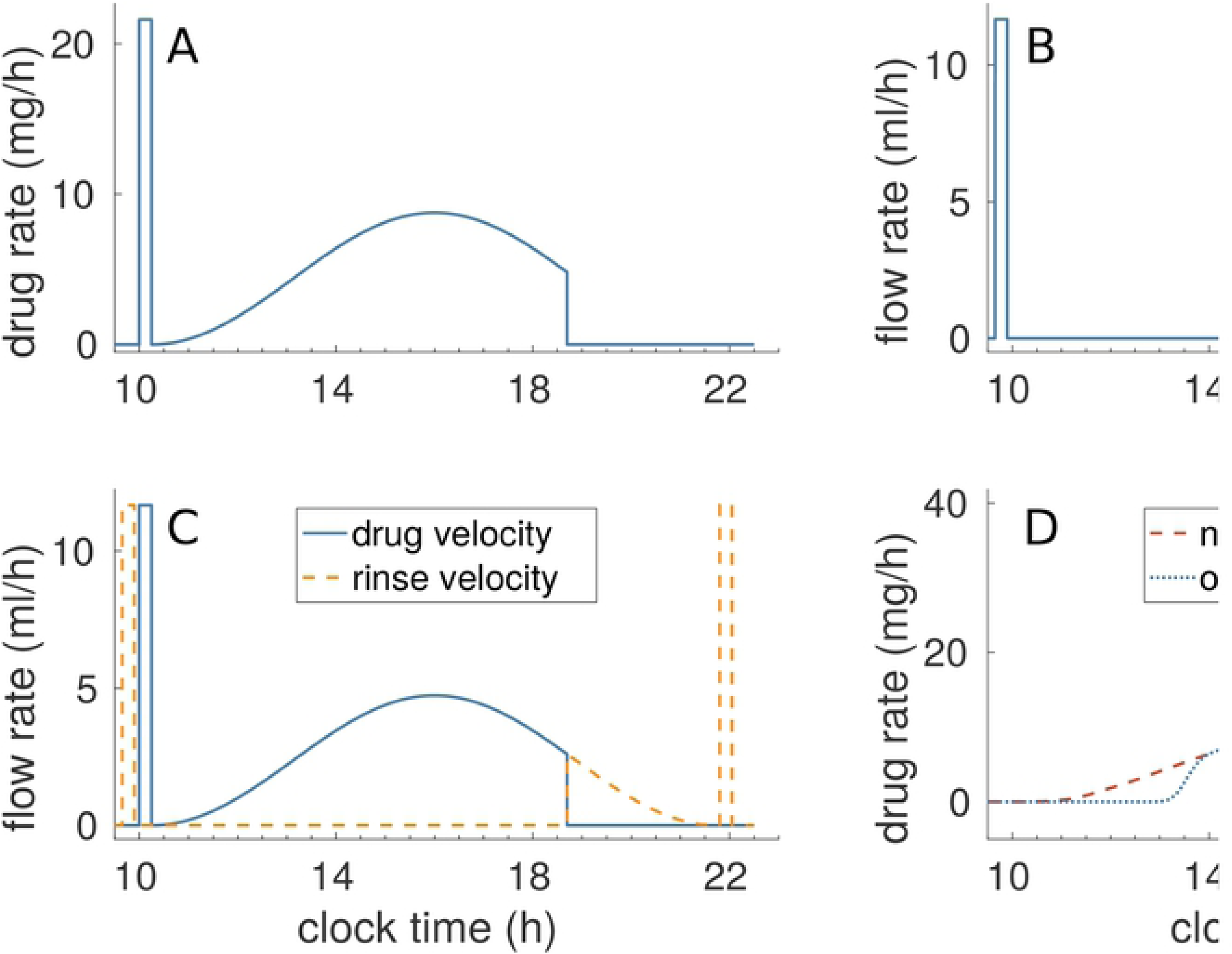
Semi-physiological model of irinotecan PK. Compartments were minimised to the most important components, Liver to accurately represent drug delivery, Blood which is measurement site and Organs to represent the rest of the body. *C*_*i*_ is the rate constant of clearance from compartment i. Irinotecan is bio-activated into its active metabolite SN38. Irinotecan was assumed to be delivered directly into the liver.

**Fig 4.**
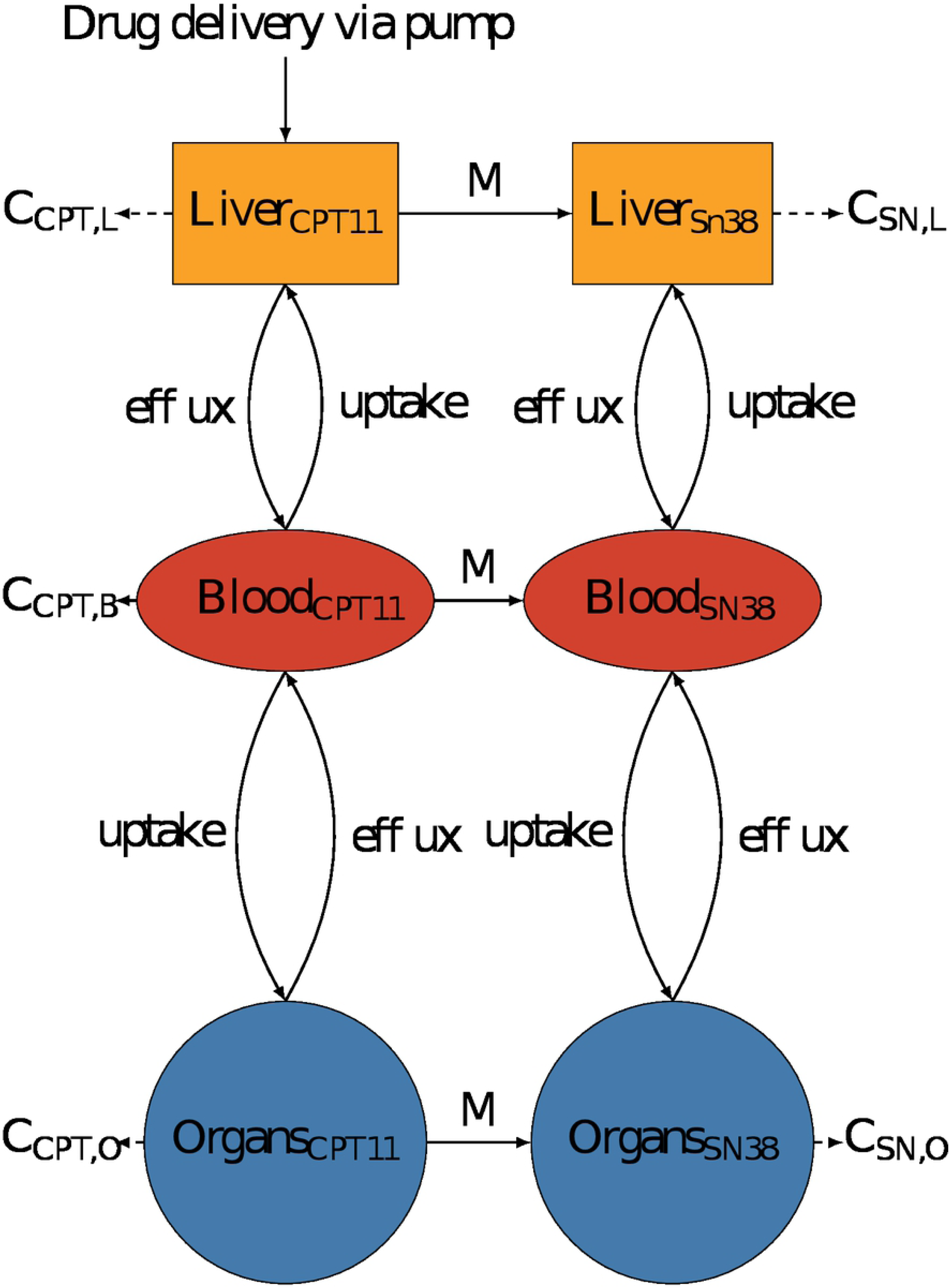
Semi-physiological model of oxaliplatin PK. Compartments were minimised to the most important components, Liver to accurately represent drug delivery, Blood which is measurement site and Organs to represent the rest of the body. *C*_*i*_ is the rate constant of clearance from compartment *i*. Each compartment contains a bound and unbound drug fraction and only unbound molecules can migrate between compartments. b and u are respectively the binding and unbinding rate constants of platinum to proteins. Oxaliplatin was assumed to be delivered directly into the liver in its unbound form.

**Fig 5.**
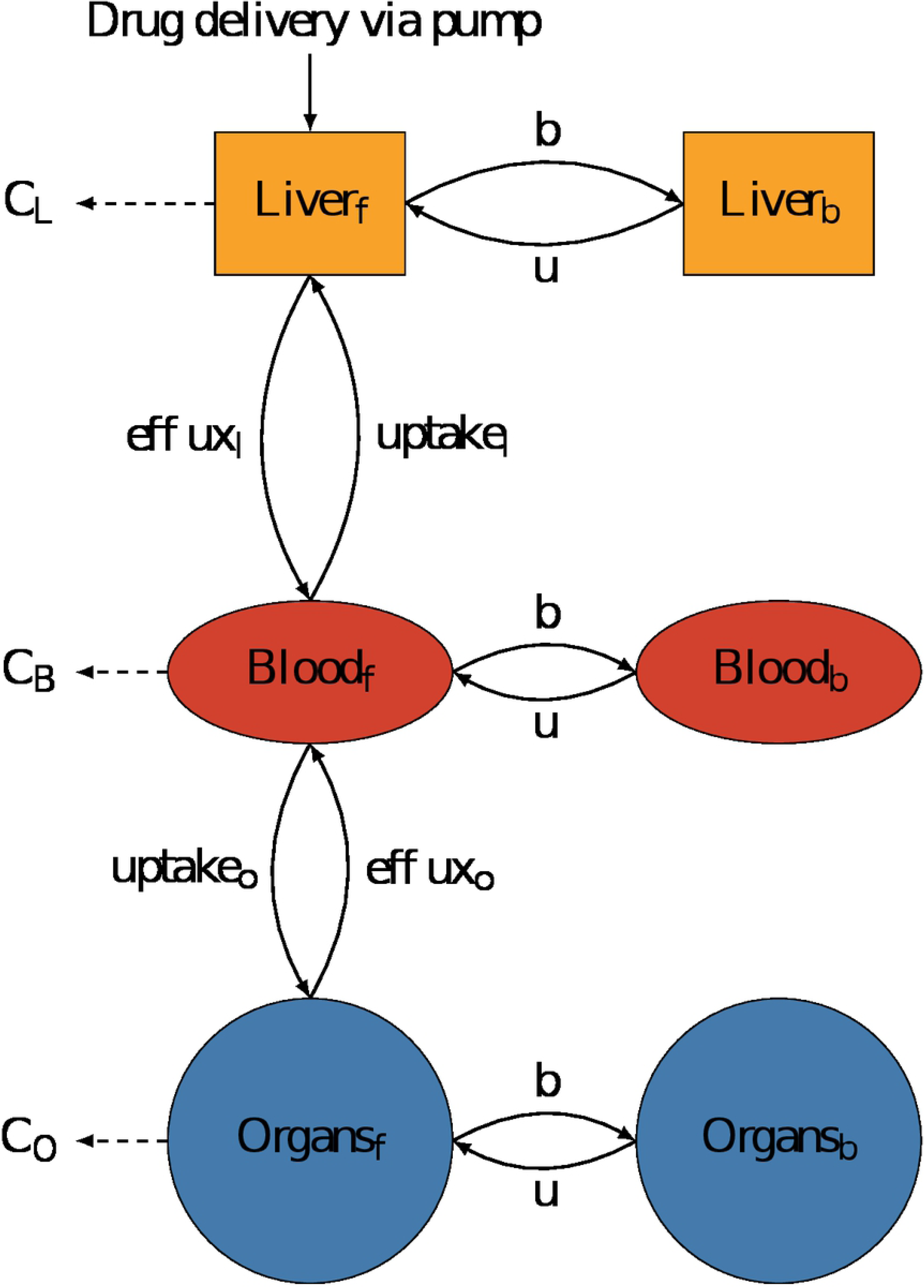
Semi-physiological model of 5-fluorouracil PK. Compartments were minimised to the most important components, Liver to accurately represent drug delivery, Blood which is measurement site and Organs to represent the rest of the body. *C*_*i*_ is the rate constant of clearance from compartment *i*. 5-fluorouracil was assumed to be delivered directly into the liver.

Parameter identifiability assessed though sensitivity analysis to cost function variations revealed poor sensitivity of the clearance rate constant in the Organs compartment for the three drugs (cf. Methods). Hence, Organs clearance was assumed to be equal to that of the Liver compartment for irinotecan and oxaliplatin and was neglected for 5-fluorouracil [24]. In the model of irinotecan and 5-fluorouracil, poor sensitivity was also obtained for transport parameters between Blood and Organs, so that Organs transport rates were assumed to be proportional to that of the Liver, and the volumes of the compartments were used to scaled parameters. Parameter likelihood profiles analysis revealed that additional constraints were needed to ensure the identifiability of all parameters (see Methods and SI). Hence, information on renal, intestinal and hepatic clearance relative rates was inferred from literature as follows. For irinotecan, CPT11 drug amount though renal clearance and though combined intestinal elimination and biliary clearance were respectively set to 34% and 51% of the total administered dose [25]. As SN38 renal elimination was documented as negligible, the metabolite was considered to only be cleared through Liver or Organs and these cleared amounts were assumed to account for 15% of the total administered dose of irinotecan [25].

Oxaliplatin clearance was set such that 55% of the total administered drug amount was cleared via the kidneys [26]. The amount of Pt bound within the Organs or within the Liver was set to 84% and 12% of the total dose, respectively [27]. 5-FU was shown to be mainly cleared through hepatic metabolism, so that the amount of drug cleared though the Liver was assumed to account for approximately 80% of the total dose [24].

The final irinotecan model had six compartments as each of the three Liver, Blood and Organs, had two sub-compartments: the parent drug irinotecan, and its active metabolite SN38 (Fig. 3). Initial irinotecan administered in the liver was assumed to be only in the form of the parent drug. Irinotecan was activated into SN38 via Michaelis Menten kinetics with the parameter estimates *K*_*m*_ taken from [28]. SN38 was considered to only be present in its bound form since the bound fraction is reported to be greater than 95% [29]. SN38 clearance terms accounted for SN38 elimination including its deactivation into SN38G though UDP-glycosyltransferases (UGTs) [28].

The oxaliplatin PK model had six compartments corresponding to bound and free platinum (Pt) molecules in the Liver, Blood and other Organs. Oxaliplatin is rapidly metabolised into platinum complex forms [26], which were not distinguished in the current data so as all metabolites of oxaliplatin were assumed to have the same PK properties in the model. Initial oxaliplatin administered in the liver was assumed to be free. Free Pt could bind to proteins and unbinding from proteins was also included in all compartments (Fig 4).

The final model for 5-fluorouracil had three compartments. The drug clearance accounted for both drug elimination and drug metabolism in each compartment (Fig 5). Protein binding of 5-fluorouracil was neglected in the model because of the low protein affinity of this drug [30]. Equations for the three models can be seen in SI.

#### Inter-patient variability in irinotecan, oxaliplatin and 5-fluorouracil PK parameters

Overall, each of the three drug models showed a very good fit to data as demonstrated by *R*^2^ values averaged over all patients of 0.86 for irinotecan, 0.79 for oxaliplatin and 0.8 for 5-fluorouracil (Fig 1, 2 and 3 and table 4, 7 and 10 in SI). These results obtained using infusion rates computed through the pump-to-patient model were compared with simulations with infusion rates equal to the profiles programmed into the pump (see SI). Using the pump-to-patient model allowed to improve SSR values by 7.9% for irinotecan, 49.5% for oxaliplatin and 12.5% for 5-fluorouracil in average for all patients, thus proving the validity of our approach. The irinotecan model had an almost perfect fit and showed a rapid accumulation of both irinotecan and SN38 in the plasma of patients (Fig 6). No obvious impact on irinotecan and SN38 plasma concentrations was observed regarding the time needed to fill the infusion tube or the 30-min glucose delivery spike, as predicted by the pump-to-patient model.

**Fig 6.**
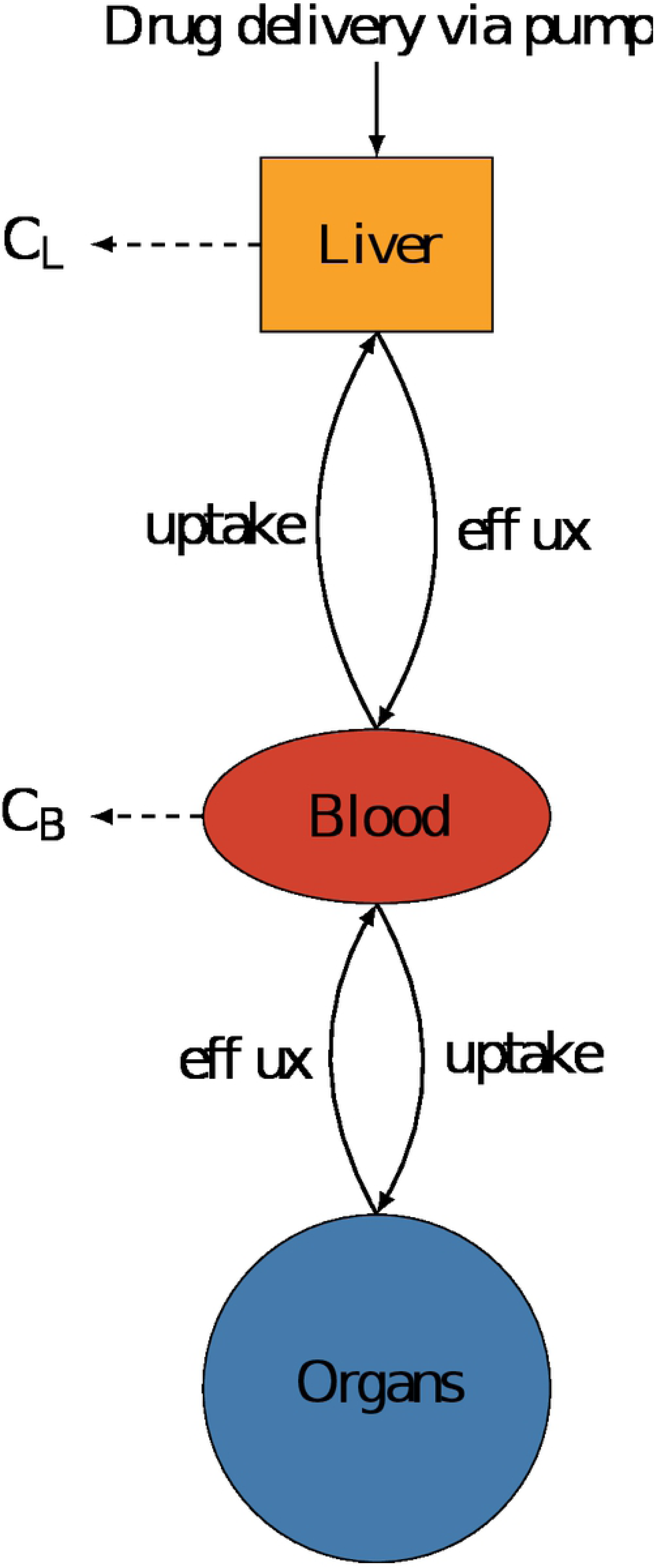
Patient data best-fit of irinotecan PK model. Each subplot represents an individual patient dataset, fit to the model independently. (a) shows the fit of irinotecan plasma concentration, (b) shows that of SN38, the active metabolite of irinotecan.

The fit for the oxaliplatin PK model captured all general trends (Fig 7). The model fit for patient 7 did not fully captured the dynamics of total Pt plasma concentration but correctly simulated free Pt concentration. The model did predict i) a delay in plasma Pt concentrations at the start of the infusion due to the pump-to-patient drug transport and ii) a spike during the glucose flush for all patients. This drug spike had an effect on the time of maximum concentration tmax of the free Pt by shifting the time by up to 6 h. The model underestimated the free platinum peak concentrations after the glucose flush for the patients with the most significant rise in concentration, that are patients 2, 3 and 7.

**Fig 7.**
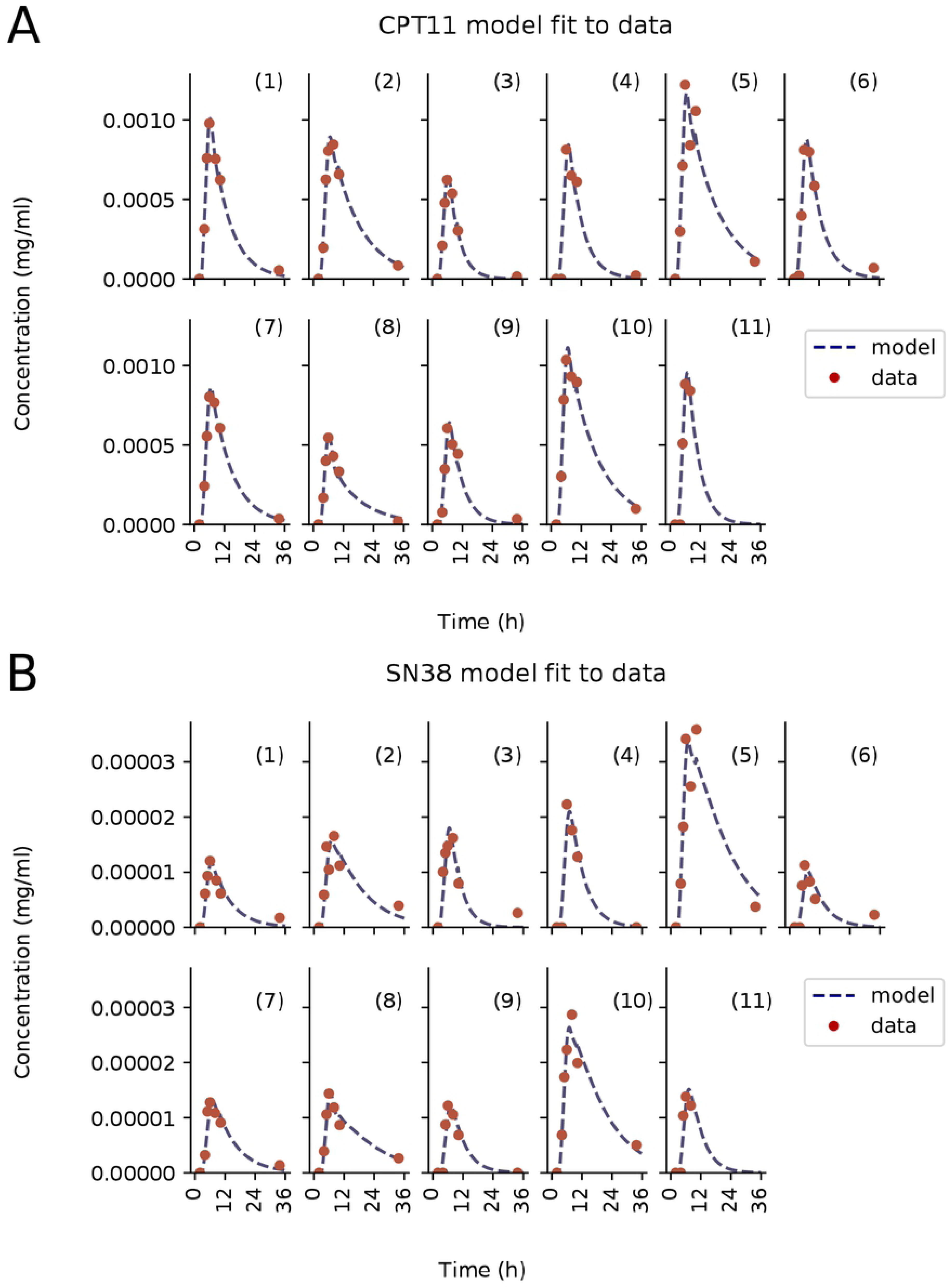
Patient data best-fit of oxaliplatin PK model. Each subplot is an individual patient data, fit to the model independently. (a) shows plasma ultrafiltrate platinum concentrations, and (b) shows plasma total platinum concentrations. PK data for Patient 11 was missing.

The 5-fluorouracil model showed a very good fit to data, despite a slight systematic under-estimation of the third datapoint in time. It predicted the glucose flush to induce a late spike in plasma drug concentration which could not be seen in the data for all patients, probably because blood sampling frequency was not high enough (Fig 8). This model-predicted spike in 5-fluorouracil concentration changed the tmax value for Patient 5, 6 and 9. The predicted spike AUC was equal to approximately 5% of the total AUC which was in agreement with the pump-to-patient model prediction. This was only calculable for 5-fluorouracil since its elimination was fast enough for its concentration to be close to zero by the time the glucose flush began.

**Fig 8.**
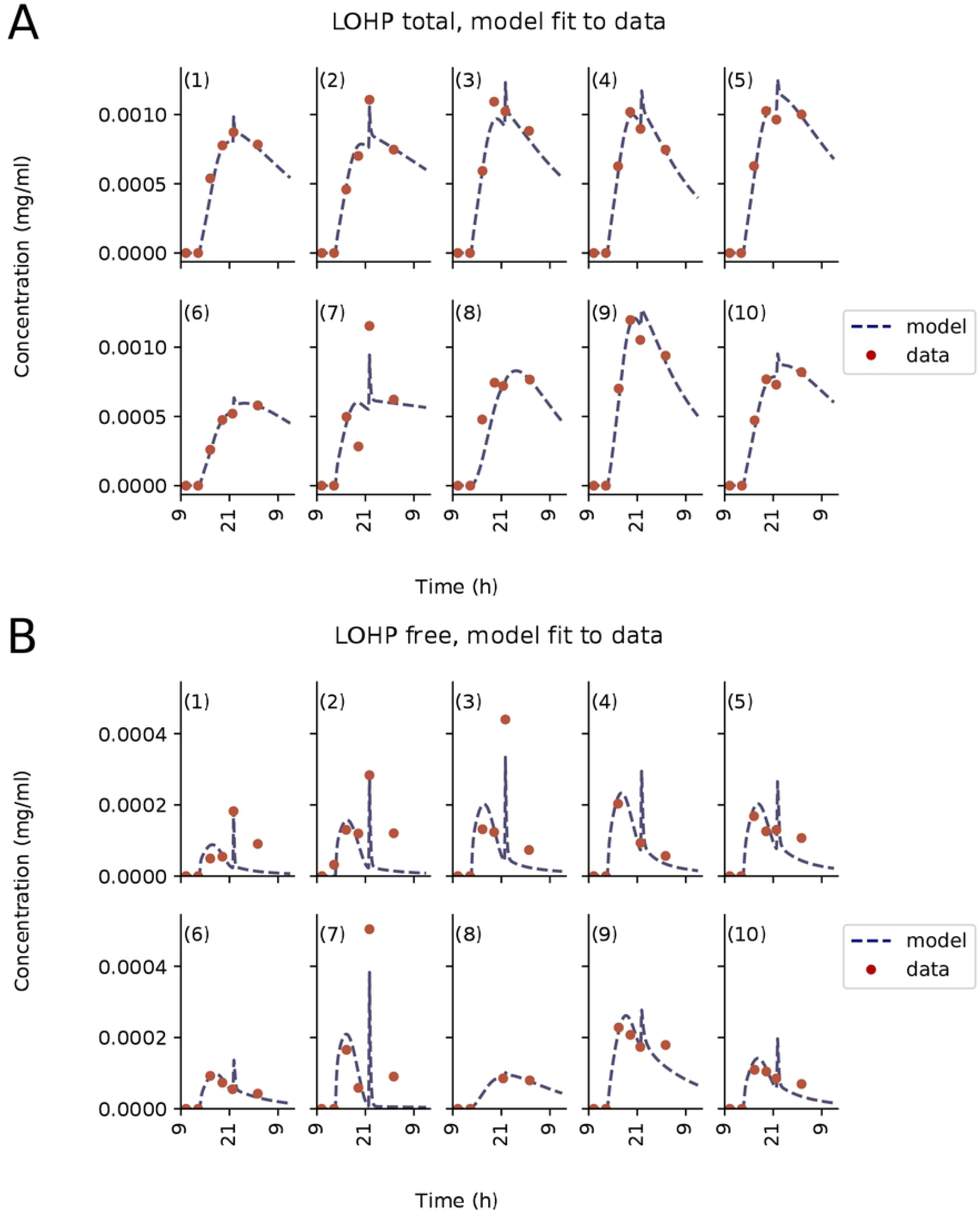
Patient data best-fit of 5-fluorouracil PK model. Each subplot is an individual patient data fit to the model independently. PK data for Patient 6 and 11 was missing.

The model fit to each individual patient PK data allowed to investigate the inter-patient variability in resulting PK parameters (Fig 9a, b, c). The CV of each PK parameter was calculated among the patient population (see SI). Interestingly, Blood-Liver and Blood-Organs transport parameters presented the highest CVs for all three drugs. Then, the mean CVs for the entire parameter set of each drug model were calculated as a single measure of inter-patient variability. Irinotecan had the smallest mean CV with a value of 73.6%, and a range from 46.1 to 170.6%. 5-fluorouracil had the second smallest mean CV at 105.69%, with the smallest range from 56.0 to 176.79%. Oxaliplatin had the largest value of mean CV, 177.87%, with the largest range from 41.15 to 302.5%. In all three models the parameters which showed the largest inter-patient variability were the uptake/efflux parameters. For each drug model, individual patient parameter sets were then utilized to identify patient clusters. The numbers of clusters were determined by minimising the validity index of Fukuyama and Sugeno *V*_*FS*_ as described in [31]. Clustering for different numbers of clusters and their respective *V*_*FS*_ can be seen in the SI (Supplementary figures 7, 8 and 9). For irinotecan, the minimum value of *V*_*FS*_ was achieved for five clusters. One cluster was composed of Patients 1, 3, 5, 7, 8, and 10, the other four patients were in a cluster on their own. The analysis for oxaliplatin concluded to two clusters, a cluster of only one patient, patient 7, and the rest of the patients being clustered together. The analysis for 5-fluorouracil revealed four clusters: 5 patients were grouped in the largest cluster (Patients 1, 2, 3, 7, and 10), two patients in the second cluster (Patients 4, 5) and the final two patients were in clusters on their own. Only patients 1, 3 and 10 were consistently clustered together for all three drugs. Once the patient PK parameters had been clustered, the mean of parameter CVs was reassessed for each cluster with 2 or more patients within. Irinotecan mean CV in the largest cluster was 47.18%, which represented a large decrease compared to the mean CV in the entire patient population equal to 73,16%. Oxaliplatin main cluster which was constituted of all patient but patient 7 had a mean CV of 165.58% as compared to 177,87% for the entire population. 5-fluorouracil’s largest cluster had a CV of 28.95% and the smaller cluster had a CV of 51.53%, which corresponded to a drastic decrease of inter-patient variability as the population mean CV was equal to 105.69%. All other clusters for each drug had only a single patient and therefore the CV could not be assessed.

**Fig 9.**
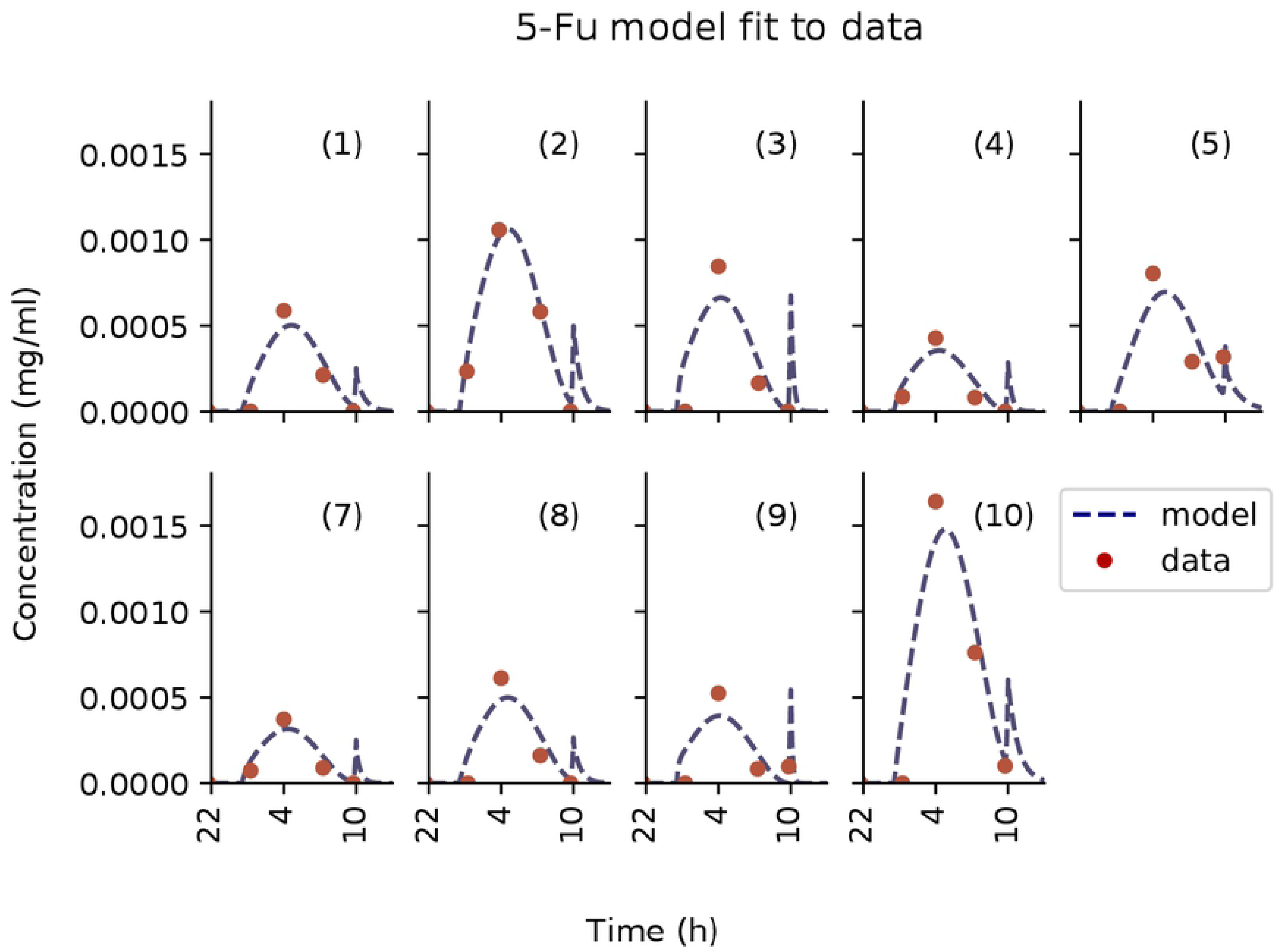
Inter-patient variability in drug PK parameters. The first line shows parameter variability across the considered patient population for irinotecan (a), oxaliplatin (b) and 5-fluorouracil (c), the colour and symbols represent the clusters each parameter set belongs to. The parameters are named with reference to the schematics of the models, the subscripts refer to the blood (B), organs (O) and liver (L). In the irinotecan parameters, additional subscripts cpt and sn refer to irinotecan and SN38 respectively. The second line shows multidimensional scaling representation of patient clustering based on their PK parameters for irinotecan (d), oxaliplatin (e) and 5-fluorouracil (f), the *x* refer to the cluster centroids and the points refer to patient PK parameters projected onto 2D plot.

## Discussion

Precision and personalized medicine requires accurate technologies for drug administration and proper systems pharmacology approaches for individual patient multidimensional data analysis. Here, plasma PK data of the OPTILIV trial in which patients received irinotecan, oxaliplatin and 5-fluorouracil through a chronomodulated schedule delivered by an infusion pump into the hepatic artery were mathematically analysed. To allow for an accurate analysis of PK patient data, a model of the pump drug delivery was successfully designed and connected to semi-mechanistic PK models. The overall framework achieved a very good fit to individual time-concentration profiles. The validity of the approach was further demonstrated by the improved data fit using the PDE explicit solution connected to PK models compared to PK models directly integrating infusion profiles that were programmed into the pump (see SI). This study gave insights into inter-patient variability and paved the path to treatment optimization.

The simulations for the pump-to-patient model showed and quantified a delay between the actual start of the pump and the time when the drug appeared in the patient blood which was due to the delay needed for the drug solution to fill up the infusion tube and eventually reach the patient. The length of this delay depends on both the drug solution concentration and the volume of the infusion tube, so that its importance was high for oxaliplatin, intermediate for 5-fluorouracil and minor for irinotecan. Temporal accuracy is key for precision medicine especially in the context of chronotherapy and chronomodulated drug delivery. Thus, the programmation of any drug administration devices need to account for these delays. The pump-to-patient model that we present here allow to adapt any infusion schemes for any drug administration devices in order to properly administer the treatment schedules initially intended by the oncologists.

In addition to such “pump-to-body” delay, the increase in free Pt concentration near 22:00 shown in the PK data was explained by a spike in oxaliplatin delivery resulting from the glucose rinse flushing out the residual oxaliplatin left within the infusion tube. This phenomenon was well captured and quantified by oxaliplatin PK model which predicted that the quantity of drug delivered in the final spike was equal to 10.7% of the total dose. The model also showed that the *t*_*m*_*ax* of oxaliplatin plasma concentration was shifted by several hours due to this delivery profile spike. In silico simulations also predicted that the glucose flush would alter the PK of 5-fluorouracil, however the sampling scheme did not cover the time when this would theoretically happen so that this prediction could not be verified experimentally. The spike only accounted for a small amount of 5-fluorouracil dose of 5.36% and may not have caused any significant detrimental effect. The delivery spike due to the glucose rinse did not seem to have influenced the plasma concentration profile of irinotecan because the drug concentration in the solution was much lower and the flow rate programmed into the pump was much higher as compared to oxaliplatin and 5-fluorouracil administration. Indeed, the spike only accounted for less than 2% of the total dose of irinotecan.

The pump-to-patient model further showed that these inconsistencies between the simulated and intended drug administration could be overcome with a simple and easily constructed adaptation of the infusion profiles, given the specific dimensions of the infusion tube. The new profile showed a much better match with the original intended administration profile.

Several published clinical studies propose mathematical models of the PK of 5-fluorouracil, oxaliplatin or irinotecan with various levels of complexity. First, a physiologically-based PK model of capecitabine, a pro-drug of 5-fluorouracil, was designed for humans [32]. However, the data available in the OPTILIV study would not allow for estimating parameters of such a detailed model. Next, numerous clinical studies have performed compartment analysis of plasma PK data from cancer patients receiving either 5-fluorouracil, oxaliplatin or irinotecan [33]. These models were designed for intravenous injection and could not be readily used for intra-arterial hepatic administration. Thus, the development of new semi-physiological PK models was necessary to include the drug delivery site as a separate compartment, that was different from the Blood compartment for which data was provided. Furthermore, the intention was also to develop more physiologically-relevant models in view of future account of circadian rhythms and vhronotherapy optimization investigations. Indeed, the developed models are called semi-physiological as the compartment volumes together with relative fractions of clearance routes were inferred from literature. These models could then be further extended to physiologically-based models by detailing the “Organ” compartment and be connected to mechanistic PD models to represent organ-specific drug PK-PD towards chrono-administration optimization.

Inter-patient differences in maximum plasma drug concentrations and in the time at which it occurred led us to further investigate variability in between subjects. Irinotecan showed the lowest mean variability. Clustering analysis indicated that patients could be classified into five clusters with respect to irinotecan PK parameters. The second largest inter-patient variability was found for 5-fluorouracil. Clustering for 5-fluorouracil showed there was four clusters. Regarding oxaliplatin, there was the largest variability between patients PK model parameters with all parameters showing high variance. Clustering according to oxaliplatin PK parameters split patients into two clusters leading to isolate patient 7. This clustering of the patients led to a reduced inter-patient variability for all drugs, especially for irinotecan and 5-fluorouracil. This decrease in CVs is not unexpected, but the significant level of reduction means this method could be used as a way to stratify patients into treatment groups with less inter-patient variability in PK profiles. The measure of inter-patient variability could be interpreted as indicators of the need for personalisation as high differences between subjects implies high potential benefit of drug administration personalisation. Here, we demonstrated that the PK of all three considered drugs displayed important inter-subject variability. The remaining clinical challenge lays in determining clinical biomarkers for stratifying patients before drug administration, in order to reach the intended plasma PK levels. In order to do so, we performed modelling analyses and identified the critical PK parameters for irinotecan, 5-fluorouracil and oxaliplatin which were the transport parameters between the Blood and either the Liver or the Organs compartments.

## Conclusion

In conclusion, a mathematical framework was designed to allow for accurate analysis of patient PK data. A model of the dynamics of the drug solution from the pump to the patient’s blood was designed, irrespective of the drug delivery device. It was used to represent the chronomodulated drug administration though the Mélodie infusion pump into the patient hepatic artery of irinotecan, oxaliplatin and 5-fluorouracil. The model revealed significant inconsistencies between the drug profiles programmed into the pump which corresponded to the drug exposure intended by clinicians and the actual plasma PK levels. Importantly, it allowed for the design of innovative drug in-fusion profiles to be programmed into the pump to precisely achieve the desired drug delivery into the patient’s blood. Next, the pump-to-patient model was connected to semi-physiological models of the PK of irinotecan, oxaliplatin and 5-fluorouracil. The overall framework achieved a very good fit to data and gave insights into inter-patient variability in the PK of each drug. Potential clinical biomarkers for treatment personalisation were suggested although further investigations in larger cohorts of patients are required. Overall, this complete framework informs on drug delivery dynamics and patient-specific PK of irinotecan, oxaliplatin and 5-fluorouracil towards precise and personalized administration of these drugs.

## Methods

### Ethics Statement

The pharmacokinetic data used in this investigation came from Lévi et al pharmacokinetic investigation [19] and the comparison study companion study of the European OPTILIV trial (ClinicalTrials.gov study ID NCT00852228), which involved nine centres in four countries [34]. The data has been analysed anonymously.

### OPTILIV clinical datasets

The OPTILIV trial included 11 colorectal cancer patients with liver metastases (7 men and 4 women with median age of 60). The combination of irinotecan, oxaliplatin and 5-fluorouracil was delivered to patients by Hepatic Artery Infusion (HAI) using the Mélodie pump [19]. The patients received an intravenous administration of cetuximab 500 mg/m2 over 2 h 30 min on the morning of day 1 which was not modelled. From day 2, chronomodulated HAI of irinotecan (180 mg/m2), oxaliplatin (85 mg/m2) and 5-fluorouracil (2800 mg/m2) were given (Fig 1). Irinotecan was delivered as a 6-h sinusoidal infusion starting at 02:00, with a peak at 05:00 on day 2. Oxaliplatin was administered as an 11h 30min sinusoidal infusion beginning at 10:15 with a peak at 16:00 on days 2, 3 and 4. 5-fluorouracil was also delivered as an 11h 30min sinusoidal infusion beginning at 22:15 with peak delivery at 04:00 at night, on days 3, 4 and 5. The superiority of this drug scheduling compared to non-circadian based administration was demonstrated for intravenous administration within several international clinical trials [15]. Between each drug infusion, there was a glucose serum flush which cleared the tubing. This was a 30-min sinusoidal infusion beginning at 09:45, and then again at 21:45 i.e. at the end of each infusion (Figure 10).

**Fig 10.**
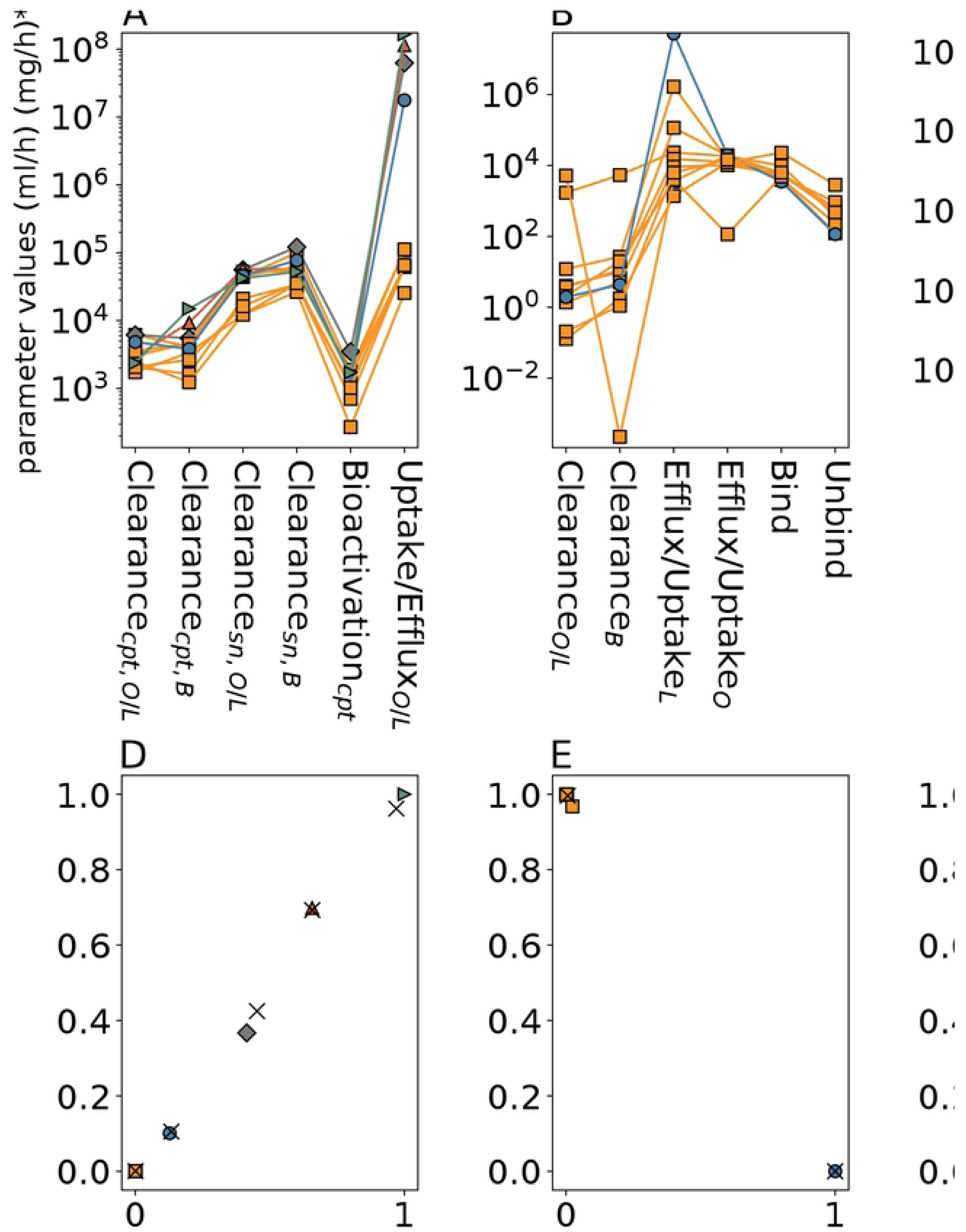
(a) Delivery profiles of irinotecan, oxaliplatin, 5-fluorouracil and glucose flushes as administered in the OPTILIV clinical trial. (b) Schematic of the Mélodie infusion pump (Axoncable, Montmirail, France) used in the OPTILIV study for hepatic artery infusion [19].

Plasma pharmacokinetics (PK) data was gathered after the first dose of irinotecan, oxaliplatin and 5-fluorouracil and measured longitudinally for each individual patient. Plasma concentrations of irinotecan and its active metabolite SN38 were determined in mg/ml at the start of infusion, then at 2, 3, 4, 6, 8 h 15 min and 31 h 45 min post HAI onset, for a total of seven time points, including baseline. Oxaliplatin concentrations were determined by measuring both platinum plasma levels, unbound and total. Oxaliplatin binds to proteins in the blood and the free Pt fraction is the biologically active one. Thus, oxaliplatin concentrations were determined at the start time of infusion, then at 3, 6, 9 h, 11 h 30 min and 17 h 15 min post HAI onset, for a total of six time points, including baseline. Plasma concentrations of 5-fluorouracil were determined at the start of infusion, then at approximately 3 h, 5 h 45 min, 9 h and 11 h 30 min post HAI, for a total of five time points, including baseline.

### Pump description

The Mélodie pump system weighs 500 g when empty (excluding drug reservoirs and batteries) and measures 160 × 98 × 34 mm. The pump consists of four channels which correspond to the four reservoirs that are connected to the pump. Each reservoir can have a maximum volume of 2 L. The four channels are controlled by four independent mechanisms which control the delivery to the infusion tube (Fig 1). For the OPTILIV study, the infusion tube comprised of two sections, the first was 135mm long with a diameter of 2.5mm, and the second section was 1500mm long with a diameter of 1mm. The two sections had a total volume of 1.84ml. The four pump reservoirs were loaded with irinotecan, oxaliplatin, 5-fluorouracil and 5% glucose solution respectively, with the latter one being used for washes in between drug infusions [35].

### Mathematical modelling

A pump-to-patient mathematical model was designed as follows, irrespective of the drug delivery device. The drug solutions dynamics from the pump to the patient’s blood was modelled using a Partial Differential Equation (PDE) considering time and 1 spatial dimension. This method was chosen as PDEs can take into account both time and space which was key for modelling systems such as pump delivery. The PDE was solved using a backward finite difference method programmed within Python 3.5.2 (https://www.python.org/). The drug PK models were based on Ordinary Differential Equations (ODEs) programmed using Python 3.5.2 and solved using the odeint function from the scipy library [36].

PK model parameter estimation involved a weighted least square approach, with conditions also placed on the drug clearance routes. The minimization of the least square cost function was performed by the Covariance Matrix adaptation Evolution Strategy (CMAES) within Python which has been shown to be successful at handling complex cost function landscapes [37]. Model goodness of fit was assessed using the sum of square residuals (SSR) and *R*^2^ values. PK model parameter numerical identifiability given the available data was investigated in a two-step process as follows. First, parameter sensitivity regarding the least-square cost function was computed via a global Sobol sensitivity analysis as a necessary condition for identifiability [38]. This method assesses the relative contributions of each parameter to the variance in the cost function obtained when parameter values are varied, and thus allows for the identification of parameters which have no effect on the cost function and are therefore not identifiable from the available dataset. This step allowed a first reduction of the PK models. Next, likelihood profiles of parameters of the reduced models were derived following the procedure outlined in [39]. Additional biological constraints derived from literature were added to ensure numerical identifiability of all parameters. This two-step model design process was undertaken as computing likelihood profiles is associated with a high computational cost.

PK models were fit to single-patient plasma PK datasets independently to obtain patient-specific parameter values. Data was available for 10 to 11 patients which was too few to undertake mixed-effect population analysis and to reliably estimate the parameters variance within a patient population [40, 41]. Sampling points at 6 hours post injection for irinotecan and 11 hours 30 mins post injection for oxaliplatin and 5-fluorouracil theoretically occurred at the same time as the start of the 30 min glucose flush, that is 9:45 for irinotecan and 5-fluorouracil, and 21:45 for oxaliplatin. As described in the results section, the flush was equivalent to the administration of the drug quantity remaining within the tube and logically influenced plasma drug concentrations. However, the exact time of patient blood collection was not reported and could vary by 10 to 15 minutes due to clinical constraints. Hence, the information of whether the blood sample was taken before or during the flush was not available. Thus, the collection time of the data points at theoretically 21:45 for oxaliplatin, 9:45 for irinotecan and 9:45 for 5-fluorouracil were unchanged if the drug concentration at the preceding data point was greater than the current one, indicating the flush might not have occurred yet. If not, the collection time was modified and set equal to the glucose peak time, which is 15min after its start time i.e. 22:00 for oxaliplatin and 10:00 for irinotecan and 5-fluorouracil, such value leading to the best model fit. Overall, the collection time was changed compared to the theoretical one for patients 1, 2, 3 and 7 for oxaliplatin, for patient 5 for 5-fluorouracil, and for no patients for irinotecan.

### Inter-patient variability and patient clustering based on PK parameters

Given the relatively small number of patients, the inter-patient variability in parameter values was assessed using a nearly unbiased estimator of coefficient of variation (CV),

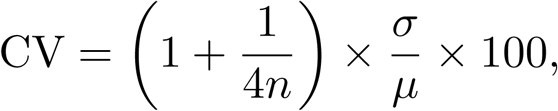

where *µ* is the parameter mean, *σ* the parameter standard deviation and n is the number of patients.

Next, fuzzy c-means clustering was used to define patient clusters based on individual PK parameters, for each drug separately. The fuzzy c-means clustering was done using a python library sckit-fuzzy (http://pythonhosted.org/scikit-fuzzy/). The method is based on the determination of cluster centroids and classification of patient parameter vectors into the clusters such that the following quantity is minimised:

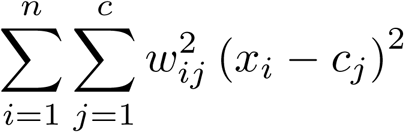

where *n*is the number of patients, *c* is the number of clusters, *x*_*i*_ is the parameter vector of patient *i, c*_*j*_ is the centroid of cluster *j. w*_*ij*_ is the probability of patient *i* belonging to cluster *j* and can be expressed as:

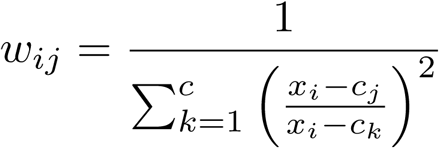

Note that, for a given patient *i*, the following holds:

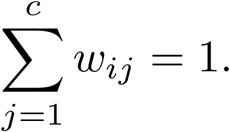

The validity function proposed by Fukuyama and Sugeno was used to determine the number of clusters for each drug. The function is defined as:

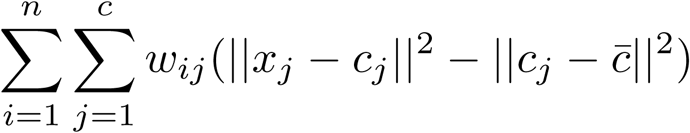

where 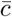 is the average of the centroids. The number of clusters were chosen between 2 and n-1 inclusively such that the VFS was minimised. Plotting the clustering results was done using a multidimensional scaling (MDS) algorithm which projects multidimensional data onto a 2D plane while keeping distance metric scaled relatively to original data (Python library sklearn.manifold [42]). Correlation coefficients between original Euclidean distance and 2D-Euclidean distance were calculated were high for all models (¿ 0.98) which showed that the MDS projections were accurate [43].

## Acknowledgements

We would like to thank Sami Al-Izzi of the University of Warwick Mathematics Institute and Dr Thomas Lepoutre (Inria, team Dracula, Lyon, France) for their discussions.

## Supporting information

**S1 Eq 1. The equations for irinotecan PK model.**

**S1 Table 1. Individual Parameter Estimates of irinotecan PK model.**

**S1 Table 2. Irinotecan PK model Parameter Mean and CV across patient population.**

**S1 Table 3. Sum of Square Residuals (SSR) for irinotecan PK model**, with either the original delivery profile, or that simulated throught the PDE pump-to-patient model. The table also shows improvement in percentages for each patient and average improvement for all patients.

**S1 Table 4.** *R*^2^ **values for irinotecan model**, with either the original delivery profile, or that simulated throught the PDE pump-to-patient model. The table also shows improvement in percentages for each patient and average improvement for all patients.

**S1 Fig. 1 Patient data best-fit of irinotecan PK model with original delivery profile** and not PDE delivery profile. Each subplot represent an individual patient dataset, fit to the model independently. The top figure shows the fit of irinotecan plasma concentration, the bottom figure shows that of SN38, the active metabolite of irinotecan. SN38 data and model simulations include both bound and free SN38.

**S1 Eq 2. The equations for oxaliplatin PK model.**

**S1 Table 5. Individual Parameter Estimates of oxaliplatin PK model**, including mean and CV of each parameter.

**S1 Table 6. Sum of Square Residuals (SSR) for oxaliplatin PK model**, with either the original delivery profile, or that simulated throught the PDE pump-to-patient model. The table also shows improvement in percentages for each patient and average improvement for all patients.

**S1 Table 7.** *R*^2^ **values for oxaliplatin model**, with either the original delivery profile, or that simulated throught the PDE pump-to-patient model. The table also shows improvement in percentages for each patient and average improvement for all patients.

**S1 Fig. 2 Patient data best-fit of oxaliplatin PK model with original delivery profile** and not PDE delivery profile. Each subplot is an individual patient data, fit to the model independently. The top figure shows plasma ultrafiltrate platinum concentrations, and the bottom figure shows plasma total platinum concentrations.

**S1 Eq 3. The equations for 5-fluorouracil PK model.**

**S1 Table 8. Individual Parameter Estimates of 5-fluorouracil PK model**, including mean and CV of each parameter.

**S1 Table 9. Sum of Square Residuals (SSR) for 5-fluorouracil PK model**, with either the original delivery profile, or that simulated throught the PDE pump-to-patient model. The table also shows improvement in percentages for each patient and average improvement for all patients.

**S1 Table 10.** *R*^2^ **values for 5-fluorouracil model**, with either the original delivery profile, or that simulated throught the PDE pump-to-patient model. The table also shows improvement in percentages for each patient and average improvement for all patients.

**S1 Fig. 3 Patient data best-fit of 5-fluorouracil PK model with original delivery profile** and not PDE delivery profile. Each subplot is an individual patient data, fit to the model independently.

**S1 Fig. 4 Parameter Identifiability for irinotecan PK model.**

**S1 Fig. 5 Parameter Identifiability for oxaliplatin PK model.**

**S1 Fig. 6 Parameter Identifiability for 5-fluorouracil PK model.**

**S1 Fig. 7 Patient parameter clustering analysis for Irinotecan.** (a) 2D vizualisation of patient clusters for different number of clusters. Centroids (stars) and patients (dots) are shown, (b) *V*_*FS*_ values for different numbers of clusters.

**S1 Fig. 8 Patient parameter clustering analysis for oxaliplatin**. (a) 2D vizualisation of patient clusters for different number of clusters. Centroids (stars) and patients (dots) are shown, (b) *V*_*FS*_ values for different numbers of clusters.

**S1 Fig. 9 Patient parameter clustering analysis for 5-fluorouracil.** (a) 2D vizualisation of patient clusters for different number of clusters. Centroids (stars) and patients (dots) are shown, (b) *V*_*FS*_ values for different numbers of clusters.

## References

1. CASyM. CASyM and the road to Systems Medicine; 2015.

2. Iyengar BR, Altman RB, Troyanskaya OG, Fitzgerald GA. Experimentation Can Enable Precision Medicine. Science. 2015;350(6258):282–283.

3. Wolkenhauer O, Auffray C, Brass O, Clairambault J, Deutsch A, Drasdo D, et al. Enabling multiscale modeling in systems medicine. Genome Medicine. 2014;6(3):4–6. doi:10.1186/gm538.

4. Anderson ARA, Quaranta V. Integrative mathematical oncology. Nature Reviews Cancer. 2008;8(3):227–234. doi:10.1038/nrc2329.

5. Agur Z, Elishmereni M, Kheifetz Y. Personalizing oncology treatments by predicting drug efficacy, side-effects, and improved therapy: Mathematics, statistics, and their integration. Wiley Interdisciplinary Reviews: Systems Biology and Medicine. 2014;6(3):239–253. doi:10.1002/wsbm.1263.

6. Boissel JP, Auffray C, Noble D, Hood L, Boissel FH. Bridging systems medicine and patient needs. CPT: Pharmacometrics and Systems Pharmacology. 2015;4(3):135–145. doi:10.1002/psp4.26.

7. Giuliano KK, Niemi C. The urgent need for innovation in I.V. infusion devices. Nursing. 2016;46(4):66–68. doi:10.1097/01.nurse.0000480617.62296.d7.

8. Hertz DL, Rae J. Pharmacogenetics of Cancer Drugs. Annual Review of Medicine. 2015;66(1):65–81. doi:10.1146/annurev-med-053013-053944.

9. Jackson SE, Chester JD. Personalised cancer medicine. International Journal of Cancer. 2015;137(2):262–266. doi:10.1002/ijc.28940.

10. Paci A, Veal G, Bardin C, Levêque D, Widmer N, Beijnen J, et al. Review of therapeutic drug monitoring of anticancer drugs part 1 - Cytotoxics. European Journal of Cancer. 2014;50(12):2010–2019. doi:10.1016/j.ejca.2014.04.014.

11. Ballesta A, Innominato PF, Dallmann R, Rand DA, Lévi FA. Systems Chronotherapeutics. Pharmacological Reviews. 2017;69(2):161–199. doi:10.1124/pr.116.013441.

12. Lévi F, Focan C, Karaboúe A, de la Valette V, Focan-Henrard D, Baron B, et al. Implications of circadian clocks for the rhythmic delivery of cancer therapeutics. Advanced Drug Delivery Reviews. 2007;59(9-10):1015–1035. doi:10.1016/j.addr.2006.11.001.

13. Dallmann R, Okyar A, Lévi F. Dosing-Time Makes the Poison: Circadian Regulation and Pharmacotherapy. Trends in Molecular Medicine. 2016;22(5):430–445. doi:10.1016/j.molmed.2016.03.004.

14. Innominato PF, Lévi FA, Bjarnason GA. Chronotherapy and the molecular clock: Clinical implications in oncology. Advanced Drug Delivery Reviews. 2010;62(9-10):979–1001. doi:10.1016/j.addr.2010.06.002.

15. Levi F, Okyar A, Dulong S, Innominato PF, Clairambault J, Lévi F. Circadian timing in cancer treatments. Annual review of pharmacology and toxicology. 2010;50(November):377–421. doi:10.1146/annurev.pharmtox.48.113006.094626.

16. Giacchetti S, Dugúe PA, Innominato PF, Bjarnason GA, Focan C, Garufi C, et al. Sex moderates circadian chemotherapy effects on survival of patients with metastatic colorectal cancer: A meta-analysis. Annals of Oncology. 2012;23(12):3110–3116. doi:10.1093/annonc/mds148.

17. Li XM, Mohammad-Djafari A, Dumitru M, Dulong S, Filipski E, Siffroi-Fernandez S, et al. A circadian clock transcription model for the personalization of cancer chronotherapy. Cancer Research. 2013;73(24):7176–7188. doi:10.1158/0008-5472.CAN-13-1528.

18. Ortiz-Tudela E, Innominato PF, Rol MA, Lévi F, Madrid JA. Relevance of internal time and circadian robustness for cancer patients. BMC Cancer. 2016;16(1). doi:10.1186/s12885-016-2319-9.

19. Lévi F, Karaboúe A, Etienne-Grimaldi MC, Paintaud G, Focan C, Innominato P, et al. Pharmacokinetics of Irinotecan, Oxaliplatin and 5-Fluorouracil During Hepatic Artery Chronomodulated Infusion: A Translational European OPTILIV Study. Clinical Pharmacokinetics. 2016; p. 1–13. doi:10.1007/s40262-016-0431-2.

20. Evans LC, Society AM. Partial Differential Equations. Graduate studies in mathematics. American Mathematical Society; 1998. Available from: https://books.google.co.uk/books?id=5Pv4LVB{_}m8AC.

21. Vauthey JN, Abdalla EK, Doherty DA, Gertsch P, Fenstermacher MJ, Loyer EM, et al. Body surface area and body weight predict total liver volume in western adults. Liver Transplantation. 2002;8(3):233–240. doi:10.1053/jlts.2002.31654.

22. Nadler SB, Hidalgo JU, Bloch T. Prediction of blood volume in normal human adults. Surgery. 1962;51(2):224–232. doi:10.5555/uri:pii:0039606062901666.

23. Sendroy J, Collison HA. Determination of human body volume from height and weight. Journal of Applied Physiology. 1966;21(1):167–172. doi:10.1152/jappl.1966.21.1.167.

24. Schilsky BRL. Biochemical and Clinical Pharmacology of 5-Fluorouracil Cellular Determinants of Sensitivity to Fluoropyrimidines. Oncology. 1998;12(7):13–18. doi:10.1053/ctrv.2002.0253.

25. Slatter JG, Schaaf LJ, Sams JP, Feenstra KL, Johnson MG, Bombardt PA, et al. Pharmacokinetics, Metabolism, and Excretion of Irinotecan (Cpt-11) following I. V. Infusion of [14C] Cpt-11 in Cancer Patients. Drug Metabolism and Disposition. 2000;28(4):423–433.

26. Graham MA, Lockwood GF, Greenslade D, Brienza S, Bayssas M, Gamelin E. Clinical Pharmacokinetics of Oxaliplatin : A Critical Review Clinical. Clinical Cancer Research. 2000;18(1):1205–1218.

27. Boughattas NA, Levi F, Fournier C, Lemaigre G, Roulon A, Hecquet B, et al. Circadian rhythm in toxicities and tissue uptake of 1,2-diamminocyclohexane(trans-1)oxalatoplatinum(II) in mice. Cancer Research. 1989;49(12):3362–3368.

28. Ballesta A, Dulong S, Abbara C, Cohen B, Okyar A, Clairambault J, et al. A combined experimental and mathematical approach for molecular-based optimization of irinotecan circadian delivery. PLoS Computational Biology. 2011;7(9):1–12. doi:10.1371/journal.pcbi.1002143.

29. Combes O, Barré J, Duché JC, Vernillet L, Archimbaud Y, Marietta MP, et al. In vitro binding and partitioning of irinotecan (CPT-11) and its metabolite, SN-38, in human blood. Investigational New Drugs. 2000;18(1):1–5. doi:10.1023/A:1006379730137.

30. Bertucci C, Ascoli G, Uccello-Barretta G, Di Bari L, Salvadori P. The binding of 5-fluorouracil to native and modified human serum albumin: UV, CD, and 1H and 19F NMR investigation. Journal of pharmaceutical and biomedical analysis. 1995;13(9):1087–93. doi:10.1016/0731-7085(95)01548-Y.

31. Wang W, Zhang Y. On fuzzy cluster validity indices. Fuzzy Sets and Systems. 2007;158(19):2095–2117. doi:10.1016/j.fss.2007.03.004.

32. Tsukamoto Y, Kato Y, Ura M, Horii I, Ishitsuka H, Kusuhara H, et al. A physiologically based pharmacokinetic analysis of capecitabine, a triple prodrug of 5-FU, in humans: The mechanism for tumor-selective accumulation of 5-FU. Pharmaceutical Research. 2001;18(8):1190–1202. doi:10.1023/A:1010939329562.

33. Deyme L, Barbolosi D, Gattacceca F. Population pharmacokinetics of FOLFIRINOX : a review of studies and parameters. Cancer Chemotherapy and Pharmacology. 2019;83(1):27–42. doi:10.1007/s00280-018-3722-5.

34. Lévi FA, Boige V, Hebbar M, Smith D, Lepére C, Focan C, et al. Conversion to resection of liver metastases from colorectal cancer with hepatic artery infusion of combined chemotherapy and systemic cetuximab in multicenter trial OPTILIV. Annals ofOncology. 2016;27(November 2015):267–274. doi:10.1093/annonc/mdv548.

35. Lévi F, Karaboúe A, Saffroy R, Desterke C, Boige V, Smith D, et al. Pharmacogenetic determinants of outcomes on triplet hepatic artery infusion and intravenous cetuximab for liver metastases from colorectal cancer (European trial OPTILIV, NCT00852228). British Journal of Cancer. 2017;117(7):965–973. doi:10.1038/bjc.2017.278.

36. Jones E, Oliphant T, Peterson P, Others. {SciPy}: Open source scientific tools for {Python}; 2001. Available from: http://www.scipy.org/{%}22.

37. Sendhoff B. Covariance Matrix Adaptation Revisited – the CMSA Evolution Strategy –. International Conference on Parallel Problem Solving from Nature. 2008;3242(Conference Paper · September 2008). doi:10.1007/b100601.

38. Zhang Xy, Trame MN, Lesko LJ, Schmidt S. Sobol Sensitivity Analysis : A Tool to Guide the Development and Evaluation of Systems Pharmacology Models. CPT: pharmacometrics & systems pharmacology. 2015;4:69–79. doi:10.1002/psp4.6.

39. Raue A, Kreutz C, Maiwald T, Bachmann J, Klingmüller U, Schilling M, et al. Structural and practical identifiability analysis of partially observed dynamical models by exploiting the profile likelihood. Bioinformatics. 2009;25(15):1923–1929. doi:10.1093/bioinformatics/btp358.

40. Kang D, Schwartz JB, Verotta D. Sample Size Computations for PK / PD Population Models. Journal of Pharmacokinetics and Pharmacodynamics. 2005;32(5-6):685–701. doi:10.1007/s10928-005-0078-3.

41. Ogungbenro K, Aarons L, Ogungbenro K, Aarons L. Sample Size / Power Calculations for Population Pharmacodynamic Experiments Involving Repeated-Count Measurements. Journal of Biopharmaceutical Statistics ISSN:. 2010;20(5):1026–1042. doi:10.1080/10543401003619205.

42. Pedregosa F, Varoquaux G, Gramfort A, Michel V, Thirion B, Grisel O, et al. Scikit-learn: Machine Learning in {P}ython. Journal of Machine Learning Research. 2011;12:2825—-2830.

43. Jaworska N, Anastasova AC. A Review of Multidimensional Scaling (MDS) and its Utility in Various Psychological Domains. Tutorials in Quantitative Methods for Psychology. 2009;5(1).

